# Evaluation of efficacy of formic acid and thermal remediation for management of *Tropilaelaps* and *Varroa* mites in central Thailand

**DOI:** 10.1101/2024.03.04.582002

**Authors:** Madison Sankovitz, Nathalie Steinhauer, Tanawat Yemor, Steven C. Cook, Jay D. Evans, Samuel D. Ramsey

**Affiliations:** Department of Ecology and Evolutionary Biology and BioFrontiers Institute, University of Colorado Boulder, CO, USA 80303; Bee Informed Partnership, Goldin Group LLC, 4500 East-West Hwy., Suite 710, Bethesda, MD, USA 2081; Department of Plant Production Technology, Rajamangala University of Technology Tawan-ok, 43 Moo 6, Sriracha, Chonburi, Thailand 20110; Agricultural Research Service, Bee Research Laboratory, United States Department of Agriculture, Beltsville, MD, USA 20705; Department of Entomology, University of Maryland, College Park, MD, USA 20742

**Keywords:** *Tropilaelaps mercedesae*, Varroa mite, western honey bee, formic acid treatment, thermal remediation, honey bee health

## Abstract

The western honey bee, *Apis mellifera*, faces a new threat from the spread of parasitic *Tropilaelaps* (Tropi) mites, specifically *T. mercedesae*, which adds additional complexity to an apicultural landscape heavily impacted by *Varroa destructor*. In this study conducted in central Thailand, we investigated the efficacy of two methods of applying formic acid and a thermal remediation technique in controlling Tropi mites and *Varroa*, focusing our attention on the reproductive stage of the mites which is restricted to capped brood cells. Results revealed that both formic acid treatments (Formic Pro and liquid formic acid) demonstrated an immediate and substantial reduction in live Tropi and *Varroa* populations, maintaining near-zero levels for the 3- week duration of the study. In contrast, thermal remediation, employing heating pads, exhibited a more gradual decline, achieving an 85.42% reduction in Tropi mites and a 92.33% reduction in *Varroa* mites by week three. Notably, heat-treated colonies experienced an unexpected resurgence in mite populations during week two. The findings contribute valuable insights into potential strategies for mitigating the threat of Tropi mites and highlight the urgency of further research to safeguard global honey bee populations.

## Introduction

The coexistence of western honey bees, *Apis mellifera* (Order: Hymenoptera), within a diverse complex of invasive pests has long created obstacles for beekeepers and raised concerns about the sustainability of these essential pollinators (Ball et al. 1988; Harrison et al. 2001; Ellis 2003; Le Conte et al. 2010; Chantawannakul et al. 2016; Chantawannakul et al. 2018).

Beekeeping is a cornerstone of the global economy and food security (Klein et al. 2007); each additional stress factor applied to this system diminishes its productivity and, subsequently, the return on investment. Honey bee colonies, with their well-organized nests, provide an ideal habitat for numerous parasites to thrive (Chantawannakul et al. 2016). Food availability year-round within the colony sustains the bees but can also support a continuous resource for parasites, promoting their proliferation. Additionally, honey bees manage a finely regulated environment within their colonies, maintaining optimal conditions that inadvertently support the growth and reproduction of parasites (Winston 1991). Moreover, highly nutritious yet defenseless brood further exacerbate the situation, as parasites often adapt to exploit this vulnerability (Jong et al. 1982).

One genus of parasites that has recently distinguished itself as a considerable threat to honey bees is *Tropilaelaps* (Tropi mites) (Ramsey 2021). Two of the four mite species in this genus have reportedly shown the capacity to adapt to the western honey bee (*T. clareae* and *T. mercedesae*) (Anderson and Morgan 2007). Native to Southeast Asia (fig. 1), *T. mercedesae* is the most concerning due to its expanding geographic range and capacity for rapid population growth (Anderson and Roberts 2013; Oštir 2014; Chantawannakul et al. 2018). Their potential expansion is foreshadowed by the now cosmopolitan distribution observed in the infamous *Varroa* mite (*Varroa destructor*) (Chapman et al. 2023). The detection of *Varroa* mites on the island of Mauritius in 2014 and Reunion Island in 2017 and the subsequent rapid expansion across these landmasses is a compelling case study, showcasing the consequences of ineffective eradication measures upon incursion of invasive honey bee parasites (Esnault et al. 2019 a, b). While data are far more limited for Tropi mites, the unchecked spread of these parasites throughout Pakistan resulting from ineffective treatment strategies is the primary factor credited with the alarming extirpation of *A. mellifera* within the country’s borders in the 1980s (Raffique et al. 2012; Khan 2020). No pesticides have been approved to date for usage against Tropi mites (de Guzman et al. 2017). These data, coupled with the ever-expanding geographic range of the Tropi mite, underscore the imperative to determine effective management measures for this invasive parasite.

**Fig. 1.**
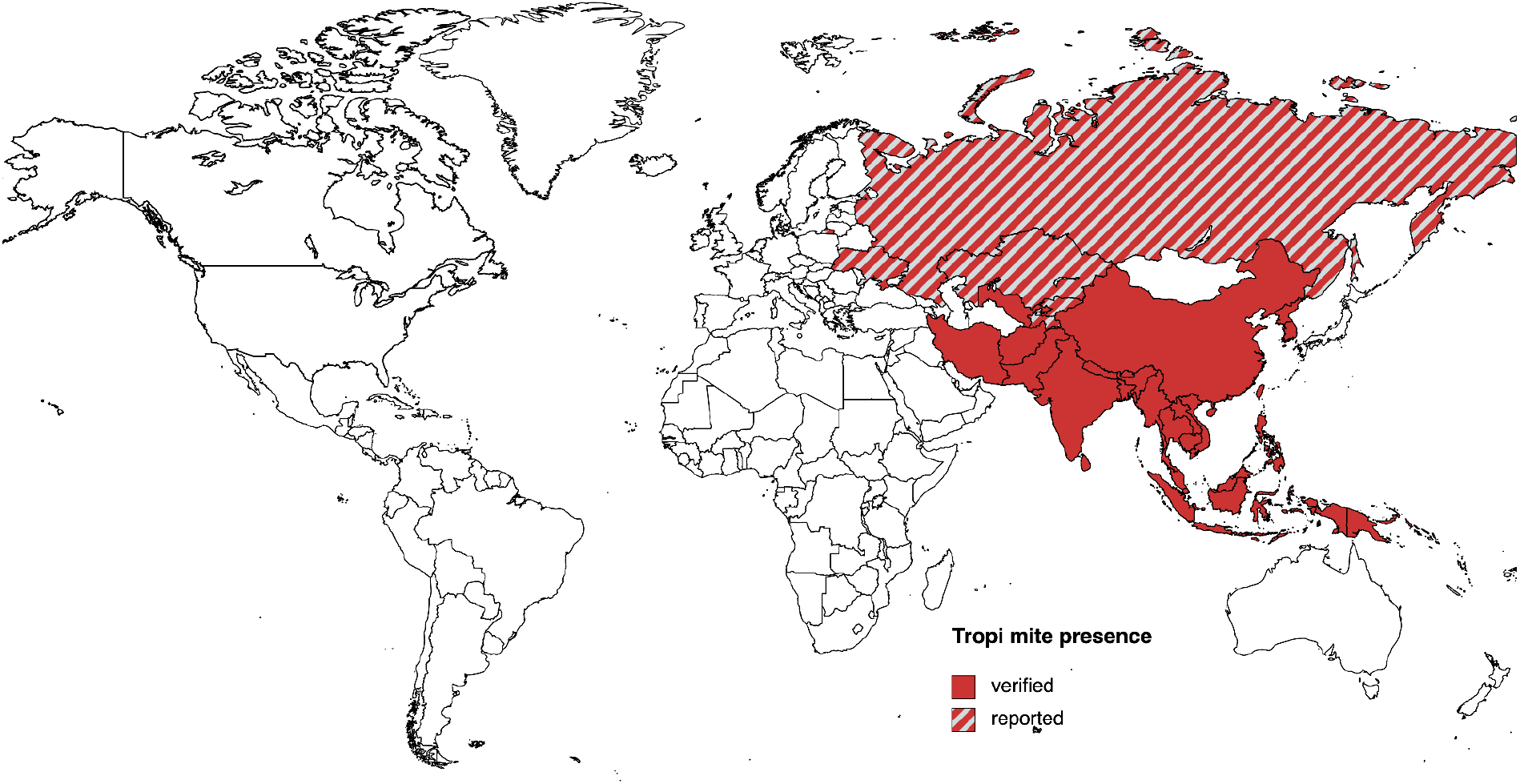
Geographic range of *Tropilaelaps mercedesae* mites. Solid red countries have verified the presence of Tropi mites, whereas striped countries have reported an infestation that has not been independently verified.

Here, we address this critical issue by investigating the efficacy of formic acid and thermal remediation techniques in controlling Tropi mites in Central Thailand. Efforts to control Tropi mites by simply repurposing treatments designed for use against *Varroa* mites without accounting for the differences between the parasites are unlikely to be successful (Pettis et al. 2017). One relevant difference in biology is the capacity of *Varroa* to exploit adult bees as a trophic resource. While *Varroa* mites feed on adult bees for several days following their reproductive phase, Tropi mites avoid this phase, using open brood as a further source of sustenance until they are ready to continue reproduction (Xie et al. 2016; Phokasem et al. 2019). As such, Tropi mites have effectively eliminated the stage at which honey bee brood parasites are the most vulnerable to chemical intervention, as the wax cell cappings that protect the brood also shield the parasites. We focused on the only registered chemical treatment for *Varroa* mites that consistently penetrates capped brood cells to kill mites sequestered with the brood. As a non-chemical measure, we chose thermal remediation for its reported capacity to impact mites within capped brood cells. For added granularity within the data set, we tested two application methods (a slow release method and a rapid release) for our chemical treatment measure.

## Methods

We conducted this research in Central Thailand in April and May of 2022 to standardize work to a single season (specifically the dry season). We selected 36 *A. mellifera* colonies from bee breeders in Kanchanaburi Province. We checked colonies for overt signs of disease (symptomatic deformed wing virus, foulbrood, etc). Strong colonies (with at least five frames of brood, substantial food stores, and no clear evidence of disease or senescence) were divided randomly and evenly between three apiaries located at Rajamangala University, Burapha University, and a Banana Tree Farm in the town of Chon Buri (Banana Garden). The treatment options were subdivided at each location by application method. Formic acid was applied via the standard method commercially available in much of the West (Formic Pro®) or via pieces of wood soaked in 85% concentrated liquid formic acid. We also tested a single method of thermal remediation using a commercially available heating pad.

In addition to the four colonies allocated to the Formic Pro, Liquid Formic, and Heating Pad respectively, each location had four control colonies, which were not treated during the study. We gave all colonies and frames unique identifying labels. Within 72 hours prior to treatment, we evaluated each colony’s mite levels via 100-cell inspection (fig. 2 & 3d). During this process, we selected a frame of capped brood from the colony for evaluation. Any frame with more than 50 capped brood cells on the front and more than 50 on the back could be chosen for this evaluation. Frames were taken back to a lab setting or well-lit outdoor area for further inspection. We opened 100 capped brood cells (half on the front and half on the back of the frame) one by one using a pair of fine forceps. We randomly chose these cells to avoid location bias in the sampling process not knowing if Tropi mites have a heretofore undescribed spatial bias for certain parts of the frame. We carefully pushed all wax composing the cap to the margins of the cell and removed the immature bee inside (fig. 3d). With the assistance of a magnifying lens and headlamp, we counted all mites within the cell. We paid particular attention to finding mites on the brood’s surface, which are similar in color and easy to overlook. Each mite egg was counted as a mite as well. We also recorded the brood’s development stage within the cell. We considered any live mites that had not started reproducing by the white-eyed pupal stage of their host (as evinced by the absence of immature mites or eggs within the cell) to be reproductive failures. Previous research has shown that mites that have not yet begun reproduction by this stage cannot reproduce successfully before their host reaches maturity (Kovacic 2017). These values established an effective baseline for mite levels and reproductive activity within the colony. Immediately after we conducted the count, we returned the frames to the colony.

**Fig. 2.**
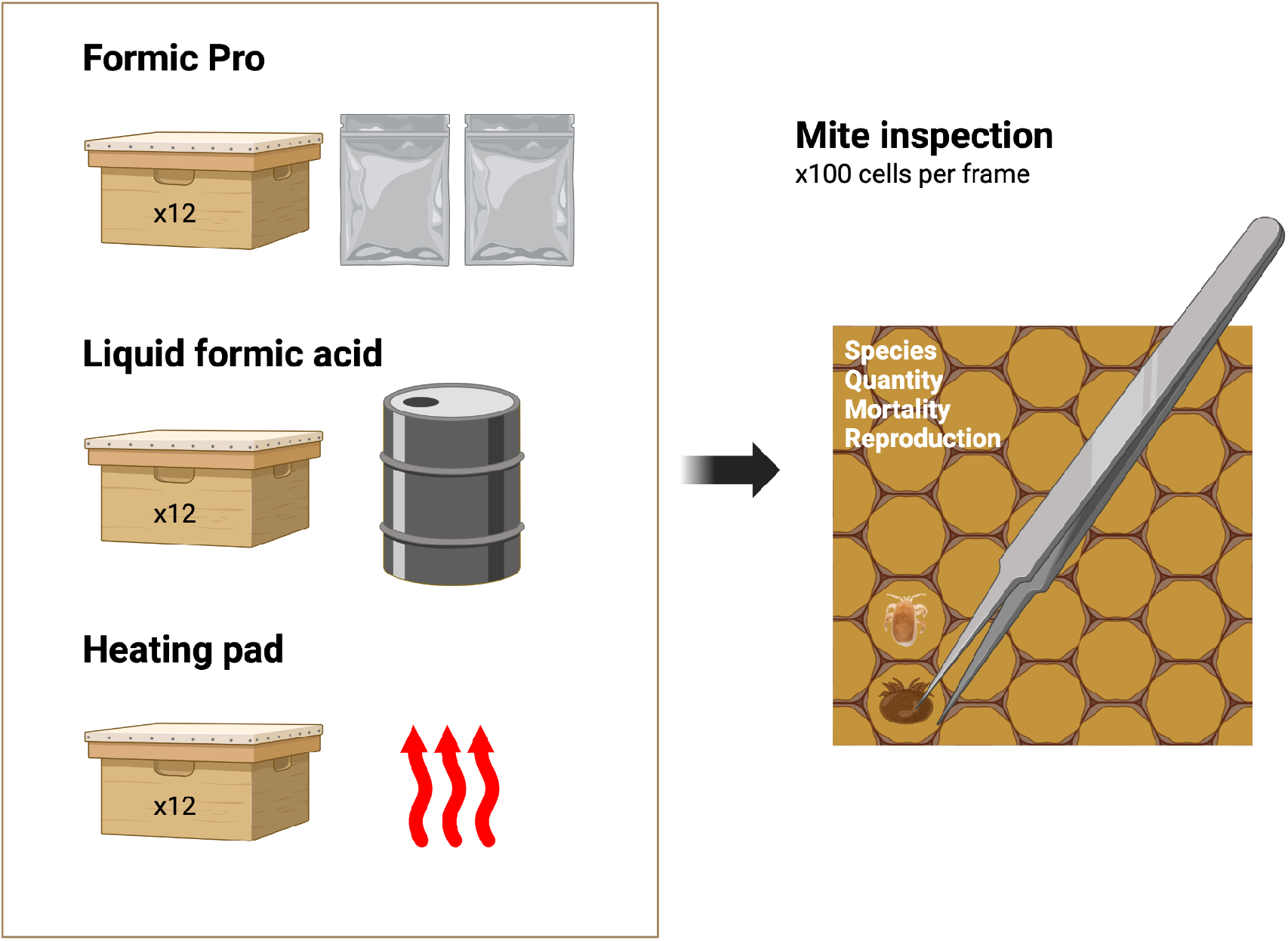
Methodology. The three treatments are shown on the left (the solar cell heat treatment was excluded due to the slow acceptance of the equipment by the bees during the trial period). We inspected 100 cells per frame weekly, identifying mite species, quantity, mortality, and reproductive success for every opened cell.

**Fig. 3.**
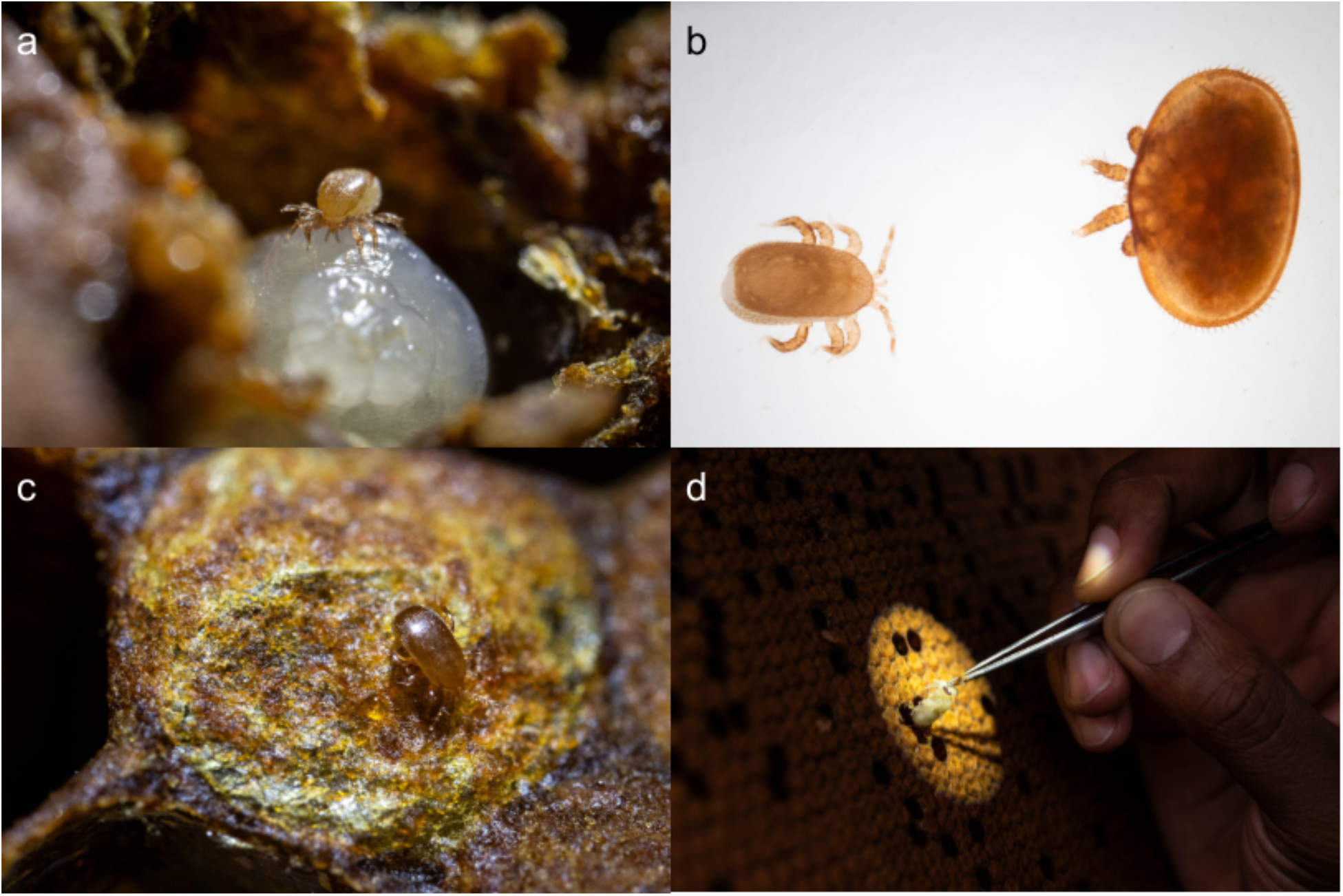
**a**. Gravid female Tropi mite (*Tropilaelaps mercedesae*) on *Apis mellifera* prepupa inside brood cell. **b**. Size comparison between *T. mercedesae* (left) and *V. destructor* (right). **c**. Tropi mite on brood cell capping. **d**. Inspection of a brood cell to determine the presence of parasitic brood mites. Observer removes the cell capping with forceps, inspects for and counts mites, removes brood and then continues the count.

Within an apiary, we exposed all colonies to their respective treatment on the same day, and we treated all three apiaries during the same week of the study. The day immediately following treatment, the first post-treatment 100-cell inspection occurred via the same procedures detailed above. Evaluation continued each successive week for three additional weeks. Another evaluation occurred at week 8, but unfortunately, repeated giant hornet attacks killed several colonies unevenly throughout our apiaries, resulting in a need to exclude the week 8 data from the final counts.

### Chemical treatment: Formic Pro

We applied Formic Pro at the most potent recommended concentration with a total dose of two sachets left in the colony for 14 days. The manufacturer recommends this extended treatment method for optimum efficacy in penetrating capped brood cells and killing the parasites within them. The sachets were placed on the tops of the frames at a staggered orientation as recommended by the manufacturer. When sampling the colony to collect brood frames for mite load evaluation, sachets were temporarily removed to access frames and returned to the colony if the treatment was still in progress. We removed the sachets for no longer than 10 minutes before being returned. After accessing the frames, we used a Draeger Gas Detector to ensure that the concentration of ambient organic acid in the colony returned to levels similar to before we opened the colony. After the prescribed 14-day treatment period, we removed the spent sachets. We also used the gas detector to compare organic acid levels between colonies twice weekly during treatment and one week after to determine that the gas had dissipated.

### Chemical treatment: liquid formic acid

We further evaluated an alternative method of applying formic acid after beekeepers in Thailand told us that they treat their colonies by applying pieces of wood soaked in formic acid beneath the frames. This particular mode of application delivers high levels of the organic acid for a shorter period than standard Formic Pro application. We could not determine the exact concentration as both chemicals were applied at a greater concentration than the Draeger Gas Detector could quantify (above 100 ppm). However, it stands to reason that the more concentrated liquid formic acid used in this application resulted in higher concentrations in the colony on the day of application than formic pro.

We obtained industrial-strength liquid formic acid from an agricultural supply store (concentration 85%). This formulation is labeled for use in, among other operations, rubber plantations to solidify liquid rubber. We cut wooden planks into pieces of 12.7 cm long and about .5 cm thick. These were soaked in formic acid for 8 hours (typically overnight) before application within the hive. On the day of application, we pushed two wooden planks per colony through the entrance of the colony positioning them parallel with the frames. Many of the bees leave the colony immediately after the organic acid impregnated planks are inserted but slowly return over the course of an hour or longer. This process was not repeated during the study period. Formic acid levels as determined by the gas detector began to decrease to levels below 60 ppm after about 48 hours post treatment. However, detectable levels of formic acid remained present in treated colonies more than a week post treatment likely due to the unventilated nature of our colonies.

### Thermal remediation: heating pad

We used Bee Hive Thermal Industries heating pads for a single heat treatment. Because of the unique structuring of the restricted hive entrances in Thai honey bee colonies, we had to first cut a wide hole into the front of the colony that would allow the insertion of the apparatus. We followed all of the manufacturer instructions using the provided insulation at the top of the colony and fastening the thermometer to the top of the centermost frame. After activating the system, it heated the colony to 41.1ºC (106ºF) for 160 minutes. After 160 minutes, the system automatically shut off, but we removed all heating pads to ensure no further heating occurred.

As the temperature within the colony increased, bees were observed leaving the colony and bearding or bundling near the front entrance. This treatment was conducted once during the treatment period.

### Thermal remediation: solar powered direct cell heating

We intended to evaluate the efficacy of an extended heat treatment measure, but the unique structuring of the equipment and time constraints prevented us from collecting enough data to effectively compare with the other treatment methods listed above. This treatment is conducted via a mode of heat delivery targeted to individual cells rather than the entire colony. Via a heating coil woven into the wax, heat is delivered through the comb so as not to heat the entire colony. This system was set to treat frames on a rolling basis over several days each week.

This treatment required us to outfit colonies with special equipment from the company Vatorex. The bees took several more weeks than expected to acclimate to the equipment, and only after six weeks did they build enough comb on the frames and begin rearing brood such that we could start the experiment. There was, however, not enough time for mites to invade the brood, and the low number of mites does not allow us to reach statistically significant conclusions about the efficacy of this treatment method.

## Results

The results of this study demonstrate the significant impact of the three different treatments, namely Formic Pro, liquid formic acid, and heat (heating pad), on the population of live Tropi and *Varroa* mites in western honey bee colonies over three weeks (fig. S1 and S2). All three treatments significantly reduced populations of both mite species by comparison to the control, with no significant difference in impact between species (fig. 4 and 5). Notably, both Formic Pro and liquid formic acid treatments exhibited immediate efficacy, rapidly reducing the number of live Tropi and *Varroa* mites. By the end of the first week post-treatment, both formic acid treatments had reduced live Tropi mite counts to zero (compared to a 56.1% increase in the control), and live *Varroa* mites were reduced by 98.78% in the Formic Pro treatment and 94.88% in the liquid formic acid treatment (compared to a 4.1% increase in the control). In contrast, the heat treatment led to a more gradual decline in live mite numbers, leading to a 85.42% reduction of Tropi and 92.33% reduction of *Varroa* mites by the end of week 3.

**Fig. 4.**
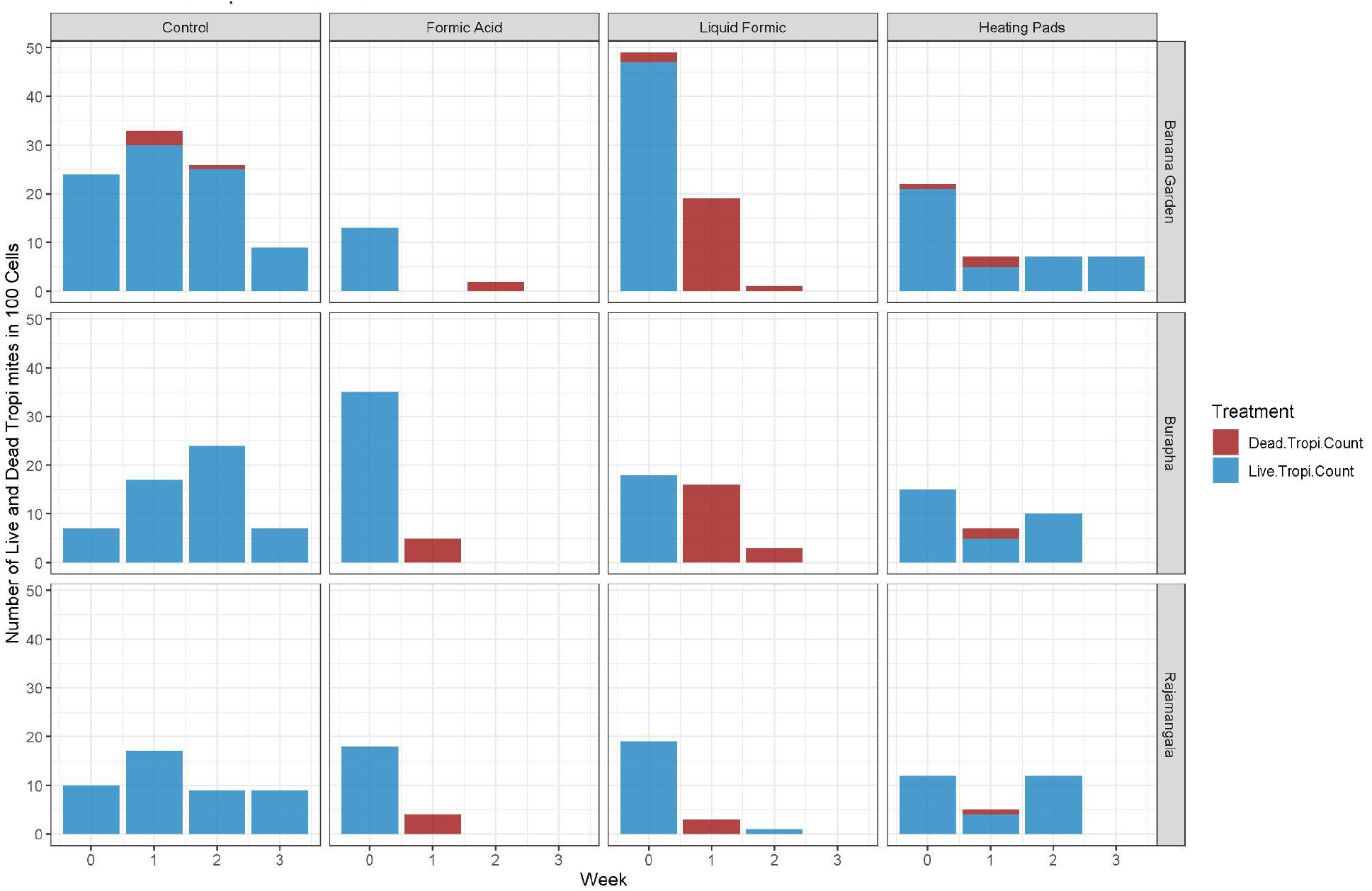
Number of live and dead Tropi mites in 100 cells per colony. All three treatments significantly reduced Tropi mite populations by comparison to the control.

**Fig. 5.**
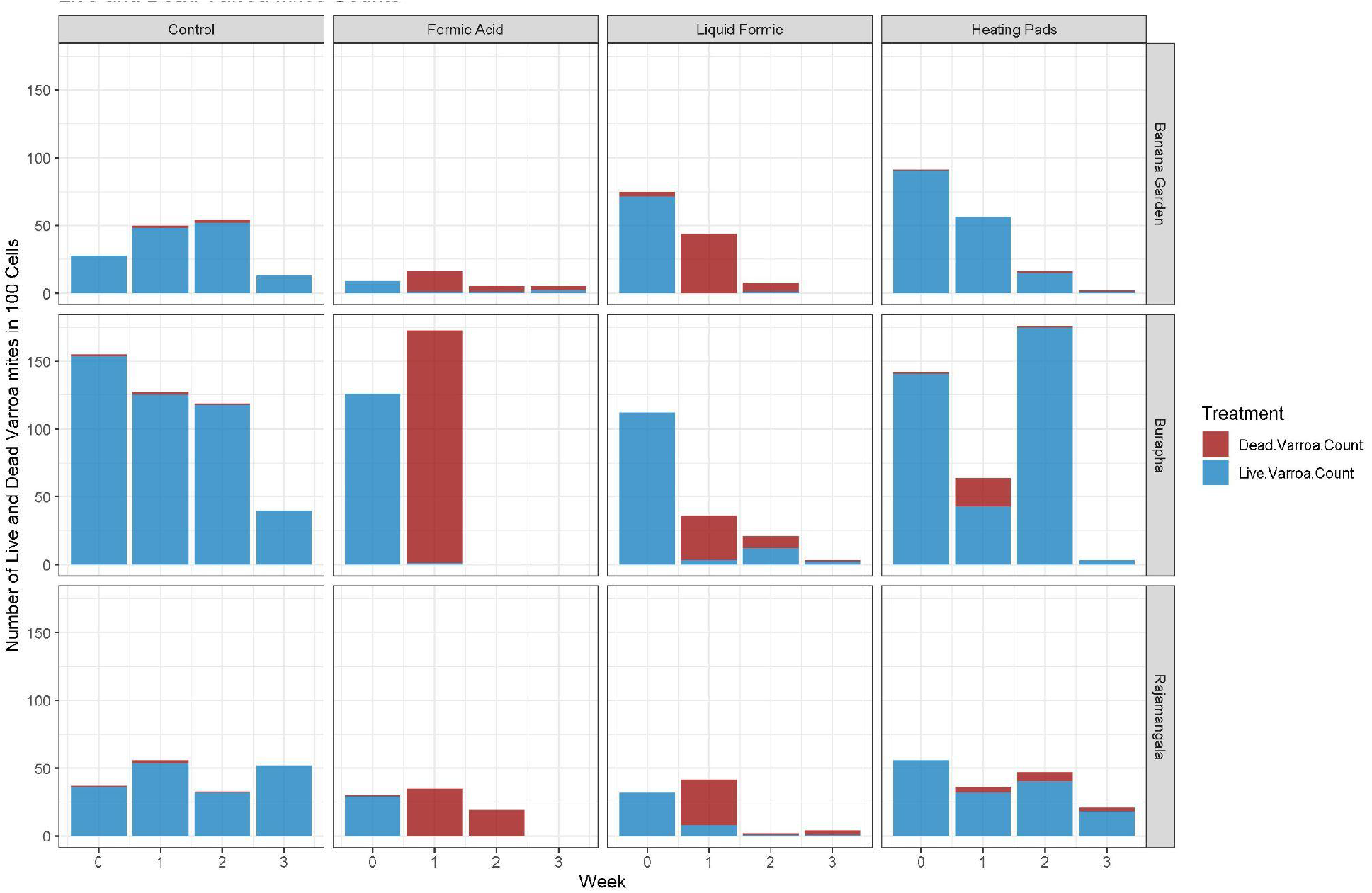
Number of live and dead *Varroa* mites in 100 cells per colony. All three treatments significantly reduced *Varroa* mite populations by comparison to the control.

Of all treatments, we observed the smallest immediate reduction in mite levels in the heat treatment. Interestingly, after a reduction (to 29.17% of live Tropi and 45.64% of live *Varroa* mite initial pre-treatment levels) between week 0 and week 1, the mite population rose (to 60.42% of live Tropi and 80.14% of live *Varroa* mite initial pre-treatment levels) in week 2. This rise in mite populations was only observed in colonies exposed to the heat treatment. This population increase was short lived as the number of Tropi mites then dropped by week 3 (to 14.58% of live Tropi and 7.67% of live *Varroa* mite initial pre-treatment levels).

Each treatment group started with an equal number of colonies but issues with frequent hornet attacks depopulated colonies unevenly. As such, our project which was originally intended to continue for an additional sampling of two weeks (at week four and week eight) had to be discontinued to ensure the values for mite levels weren’t impacted by hornet-related colony mortality.

The occurrence of dead mites (distinguished from the absence of live mites) varied among treatments. The largest impact of formic acid treatments and the only direct impact of the heat treatment were expected in week 1. There were far more dead Tropi mites in the cells of the liquid formic acid treatment (44.19% of the original mite population detected at the outset of the study) than those of the Formic Pro treatment (13.64% of the original mite population detected at the outset of the study) during week 1. Conversely, there were fewer *Varroa* mites in the cells of the liquid formic acid treatment (50.68% of the original mite population detected at the outset of the study) than those of the Formic Pro treatment (134.54% of the original mite population detected at the outset of the study) during week 1. Overall, the amount of dead mites resulting from formic acid treatments was much greater than the heat treatment. There was no practical difference between dead Tropi and *Varroa* mites found during week 1 in cells that received the heat treatment (10.20% of all Tropi and 8.65% of all *Varroa* mites detected at the outset of the study).

More *Varroa* survived each of the 3 treatments than Tropi. Notably, while we found clear evidence of dead *Varroa* mites on the heating pad post-treatment, no dead Tropi mites were observed, even when dead mites were present within the colony.

Co-infections, cells containing both Tropi and *Varroa* mites, were present in every treatment group at week 0 (fig. S3). There was no notable variation in reproductive success across treatments nor weeks; it stayed at or below an approximate 50% success rate (fig. S4 and S5).

## Discussion

All treatment strategies in our study produced a noteworthy reduction in both Tropi and *Varroa* mite populations. The formic acid treatments exhibited an immediate and remarkable decline in live mites to near-zero levels, while the heat treatment also induced an immediate reduction, though to a lesser extent. Intriguingly, colonies treated with formic acid maintained an effectively eradicated mite population throughout the trial, in contrast to the heat-treated colonies, which experienced a resurgence in week 2. This resurgence may be linked to the heat treatment stimulating mite reproduction, similar to what has been found in *Tetranychus cinnabarinus*, another mite that is an agricultural pest (Chen et al. 2023). Alternatively, the single application of heat treatment at the trial’s commencement, coupled with incomplete population eradication, may have allowed the mites to regrow from a base population while no longer under the influence of the treatment (unlike the formic acid treatments, which maintained measurable levels for more than a week with liquid formic and more than 2 weeks with Formic Pro).

Notably, the number of dead mites resulting from formic acid treatments greatly surpassed that from the heat treatment. Further, there was a pronounced difference in the number of dead mites observed between formic treatments with Formic Pro having substantially fewer dead mites. This discrepancy may stem from a difference in how the different formic acid application methods induce bees to remove brood. Both applications for formic acid achieved substantial mite reduction with no significant difference between them so finding fewer dead mites below the cell capping likely isn’t the result of a difference in efficacy of the two methods. It may, however, be the case that the bees are more inclined to uncap cells and remove brood and dead mites when the colony is exposed to the extended release of organic acid with Formic Pro rather than with the more rapid release of liquid formic.

While we observed clear evidence of dead *Varroa* mites on the heating pad post-treatment, no dead Tropi mites were found on the heating pad, even when dead mites were present within capped brood cells. This observation could be a result of Tropi mites spending limited time outside capped brood cells (a characteristic behavior of their life cycle), potentially trapping dead mites within the cells they parasitize (de Guzman et al. 2017).

Observations of adult bee mortality were noted in both formic acid application methods and, to a far lesser extent, in heat-treated colonies. We can likely attribute this mortality to the unventilated nature of the colonies. Best practices for using formic acid to treat bees include removing entrance reducers to improve ventilation (NOD Apiary Products 2018a,b). Many hives in Thailand have permanently reduced entrances, designed to restrict hornet access and this unintentional reduced ventilation allowed organic acid concentrations to exceed 100 ppm.

Additionally, finding a stretch of three or more days below 29.5ºC in Thailand for chemical application proved challenging, potentially contributing to variations in post-treatment bee loss.

Despite these challenges, the widespread use of Formic Pro in the Western world suggests that formic acid application could achieve brood parasite mortality with minimal impact on adult bees. However, successful application requires more ventilation, achieved through fully opened hive entrances, and adherence to prescribed temperature recommendations (NOD Apiary Products 2018a,b). These guidelines are critical when working with this chemical, whether using the Formic Pro method or liquid formic acid application. While liquid formic acid application shows promise as an effective treatment measure, Formic Pro treatment leads to lower levels of bee mortality. Liquid formic as a less standardized system resulted in unpredictable mortality levels potentially resulting from the different varieties of wood used. We standardized for the size of the wood and how long to soak it in formic but not the kind of wood which can impact porosity and thus absorption and release of the organic acid. With further refinement, this method may offer the best blend of accessibility, cost-effectiveness, and efficacy of the options available in much of the world. As shown here, heat treatments are possible management measures but need extensive testing to reduce variability and increase their confidence interval.

The *Varroa* mite (a relative of Tropi) is one of the most economically costly invasive species globally, underscoring the value of investing attention in understanding Tropi mites while the potential for eradication or preventing establishment is still possible (Nentwig and Vaes-Petignat 2014). As things currently stand, Tropi mites have yet to invade North or South America but they have become invasive in more than half a dozen countries to date. These data can help inform treatment efforts where these parasites are now established and eradication attempts in regions where they have not yet established populations.

Tropi mites, smaller and more agile than their *Varroa* counterparts, are a particularly concerning threat in the world of beekeeping for multiple reasons (de Guzman et al. 2017). About one-third the width of *Varroa* mites, these tiny pests inflict multiple feeding wounds on the pupal stage of bees (Anderson and Morgan 2007), unlike *Varroa* mites, which only makes a single wound. This behavior can lead to permanent deformities in the developing bees. Even more alarming, Tropi mites have been found to spread Deformed Wing Virus and Acute Bee Paralysis Virus, and there is a possibility that they might also transmit other viruses such as Black Queen Cell Virus (Forsgren et al. 2009; Chanpanitkitchote et al. 2018). Tropi mites exhibit faster population growth than *Varroa* mites, with a brief phoretic phase and relatively more time spent in the reproductive phase (de Guzman et al. 2017). Once a colony is infested, it can rapidly collapse.

The addition of new pests such as Tropi mites to a crowded stage of stress factors, which already includes unpredictable weather patterns driven by climate change, *Varroa* mites and their associated viral complex, food products tainted with insecticide residues, and inadequate nutrition, may likely be devastating (Currie et al. 2010; Chantawannakul et al. 2018). Stress factors that spread like contagions, as do Tropi mites, within an apiary often have the most significant impact on the health of honey bee populations because they link the fate of a single colony to any nearby. Densely packed apiaries lend themselves to outbreaks threatening the most lucrative apicultural operations, commercial apiaries. Moreover, Tropi mites rapidly move colonies towards collapse, reducing the net value of the colony itself to zero (Chantawannakul et al. 2016). More concerning still, is the generalism shown in Tropi as it suggests a potential toattack other bee species outside the genus *Apis*. Should reports of Tropi mites on bumble bees and carpenter bees be independently verified, Tropi will officially be considered a danger to more than just apiculture but also native pollinator populations (many of which are already beleaguered; Abrol & Putatunda 1996, Putatunda & Abrol 2003). To protect bee populations worldwide, further investigation of these Tropi mites and their impact on bee health and survival is urgently needed.

## Competing interests

The authors have no relevant financial or non-financial interests to disclose.

## Notes

### Competing Interest Statement

The authors have declared no competing interest.

