## Supplementary information for "Evaluation of efficacy of formic acid and thermal remediation for management of *Tropilaelaps* and *Varroa* mites in central Thailand"

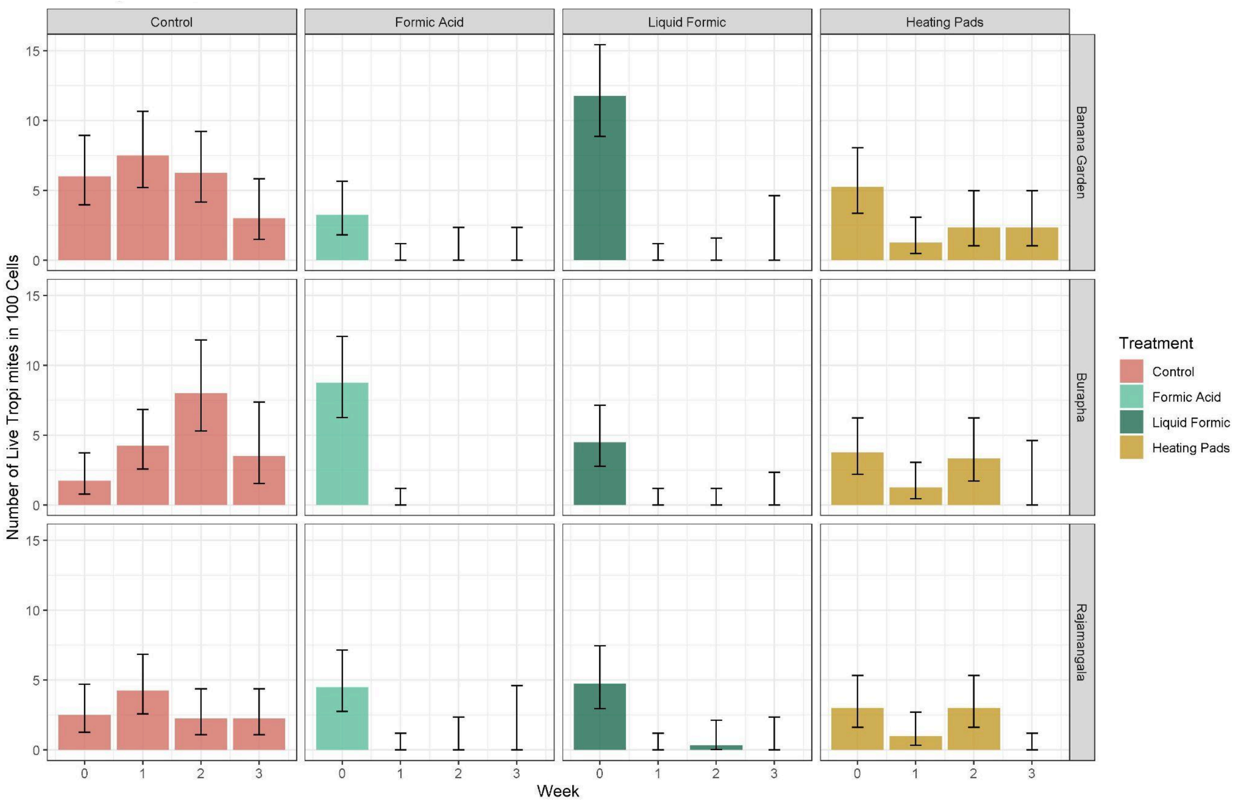


Figure S1. Number of live Tropi mites in 100 cells per colony over three weeks, separated by treatment. All three treatments significantly reduced populations of Tropi mites by comparison to the control. Notably, both Formic Pro and liquid formic acid treatments exhibited immediate efficacy, rapidly reducing the number of live Tropi mites. By the end of the first week post-treatment, both formic acid treatments had reduced live Tropi mite counts to zero. In contrast, the heat treatment led to a more gradual decline in live mite numbers, leading to an 85.42% reduction of Tropi by the end of week 3.

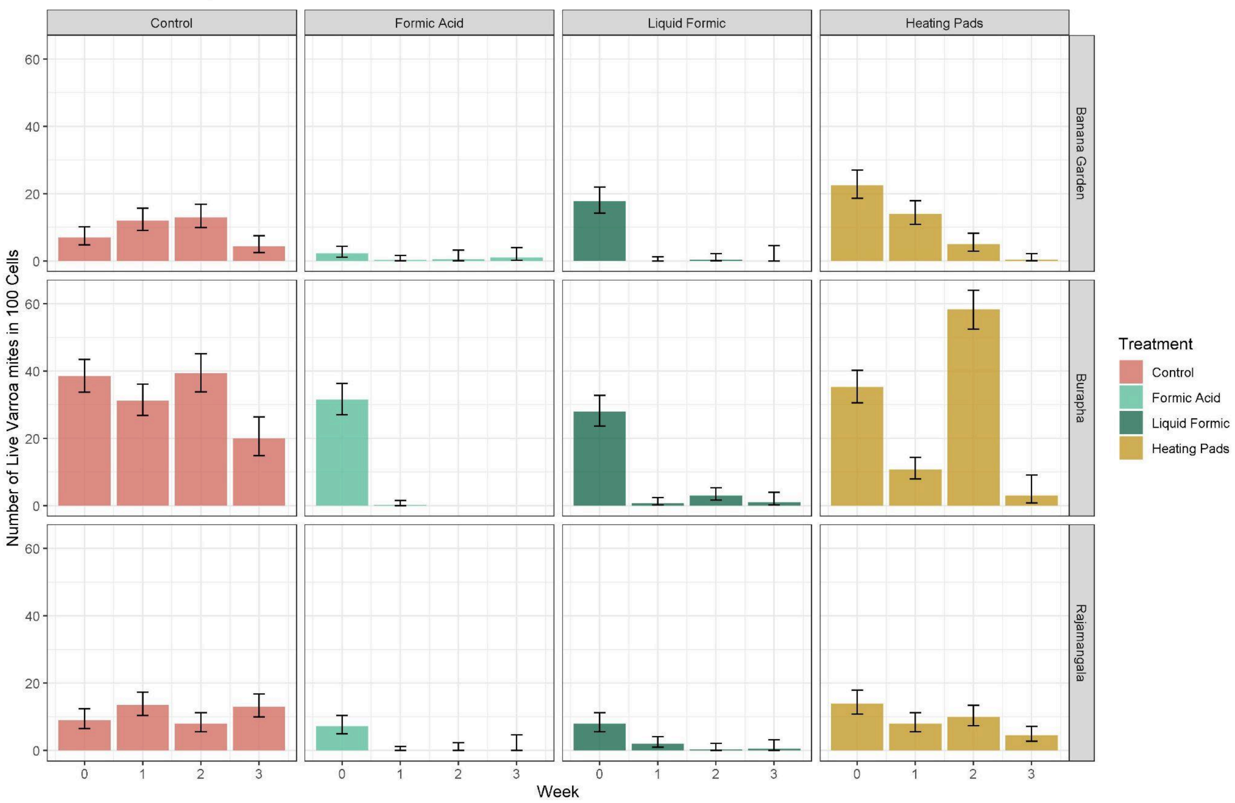


Figure S2. Number of live *Varroa* mites in 100 cells per colony over three weeks, separated by treatment. All three treatments significantly reduced populations of *Varroa* mites by comparison to the control. Notably, both Formic Pro and liquid formic acid treatments exhibited immediate efficacy, rapidly reducing the number of live *Varroa* mites. By the end of the first week post-treatment, *Varroa* mites were reduced by 98.7805% in the Formic Pro treatment and 94.8837% in the liquid formic acid treatment. In contrast, the heat treatment led to a more gradual decline in live mite numbers, leading to a 92.33% reduction of *Varroa* mites by the end of week 3.

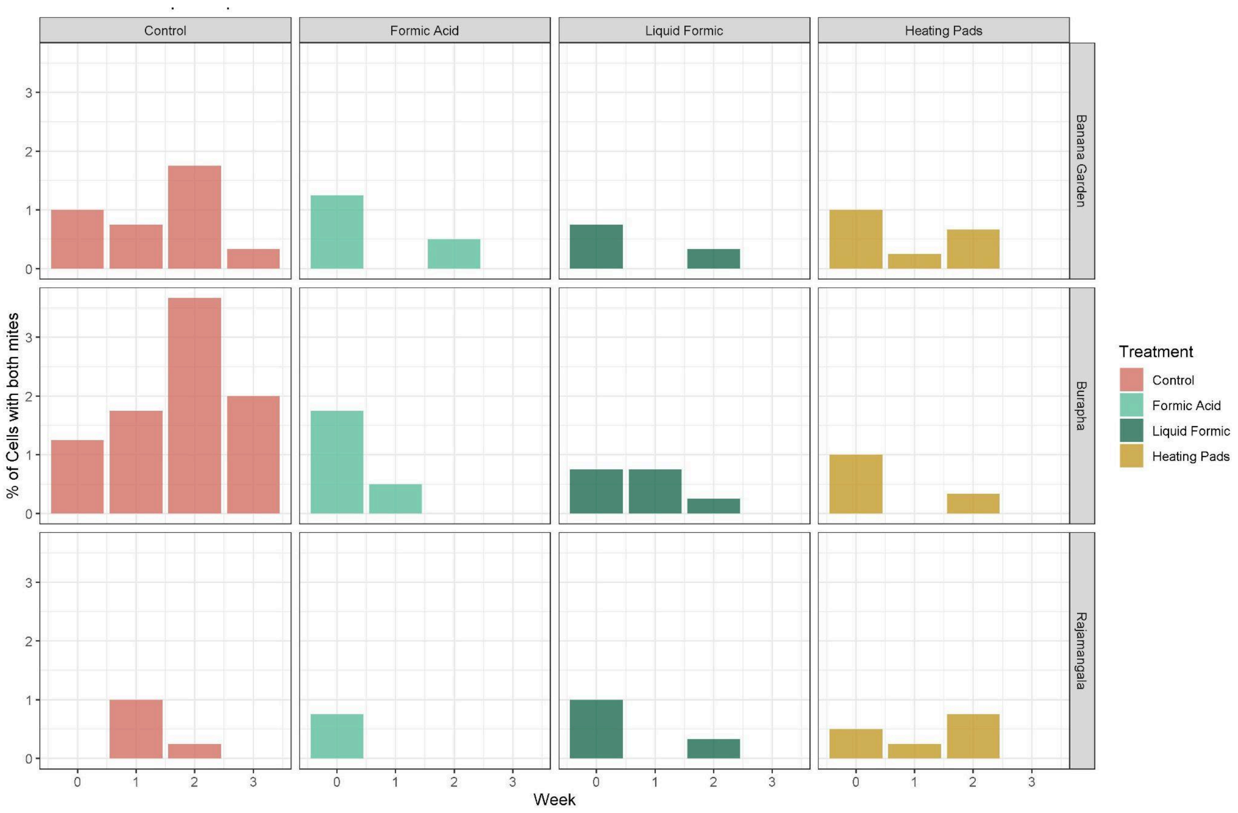


Figure S3. Percentage of cells containing both Tropi and *Varroa* mites. Co-infections were present in every treatment group at week 0.

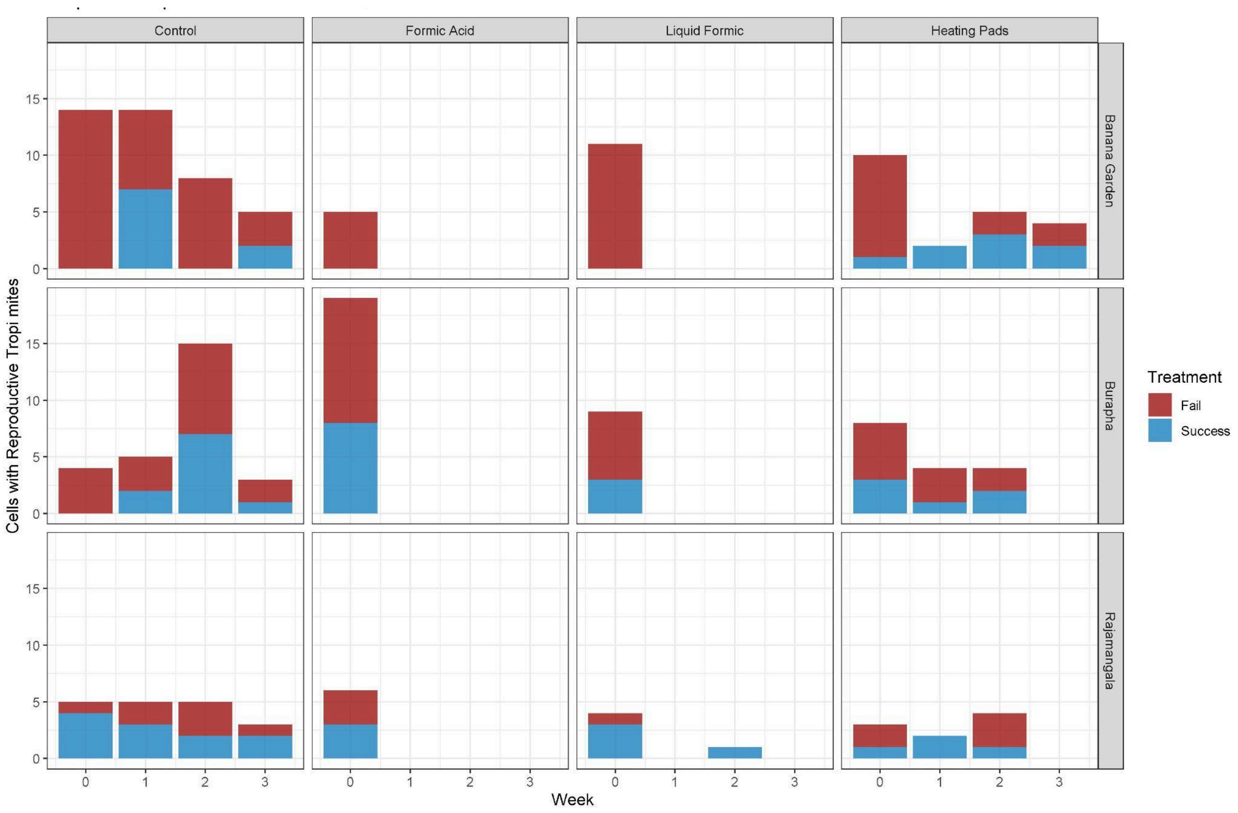


Figure S4. Cells containing reproductive Tropi mites based on treatment success across three trial weeks. There was no notable variation in reproductive success across treatments or weeks; it stayed at or below an approximate 50% success rate.

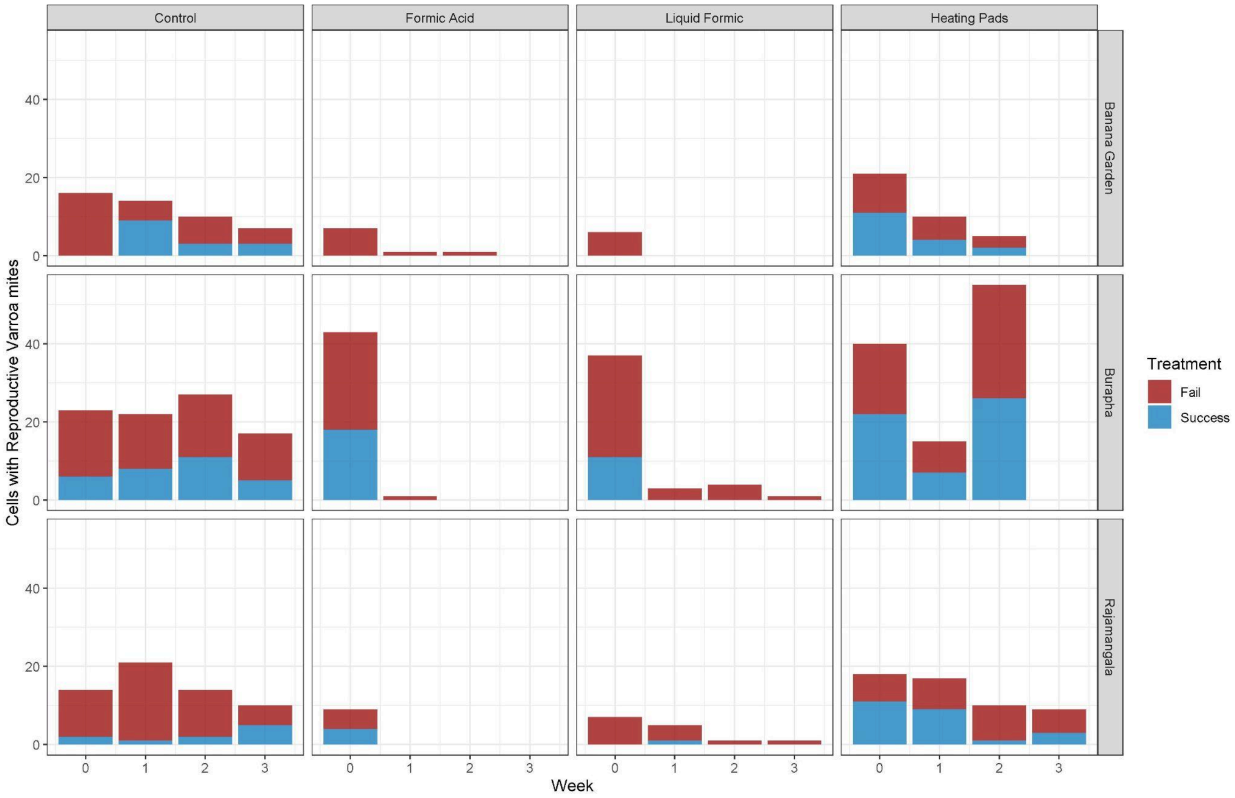


Figure S5. Cells containing reproductive *Varroa* mites based on treatment success across three trial weeks. There was no notable variation in reproductive success across treatments or weeks; it stayed at or below an approximate 50% success rate.

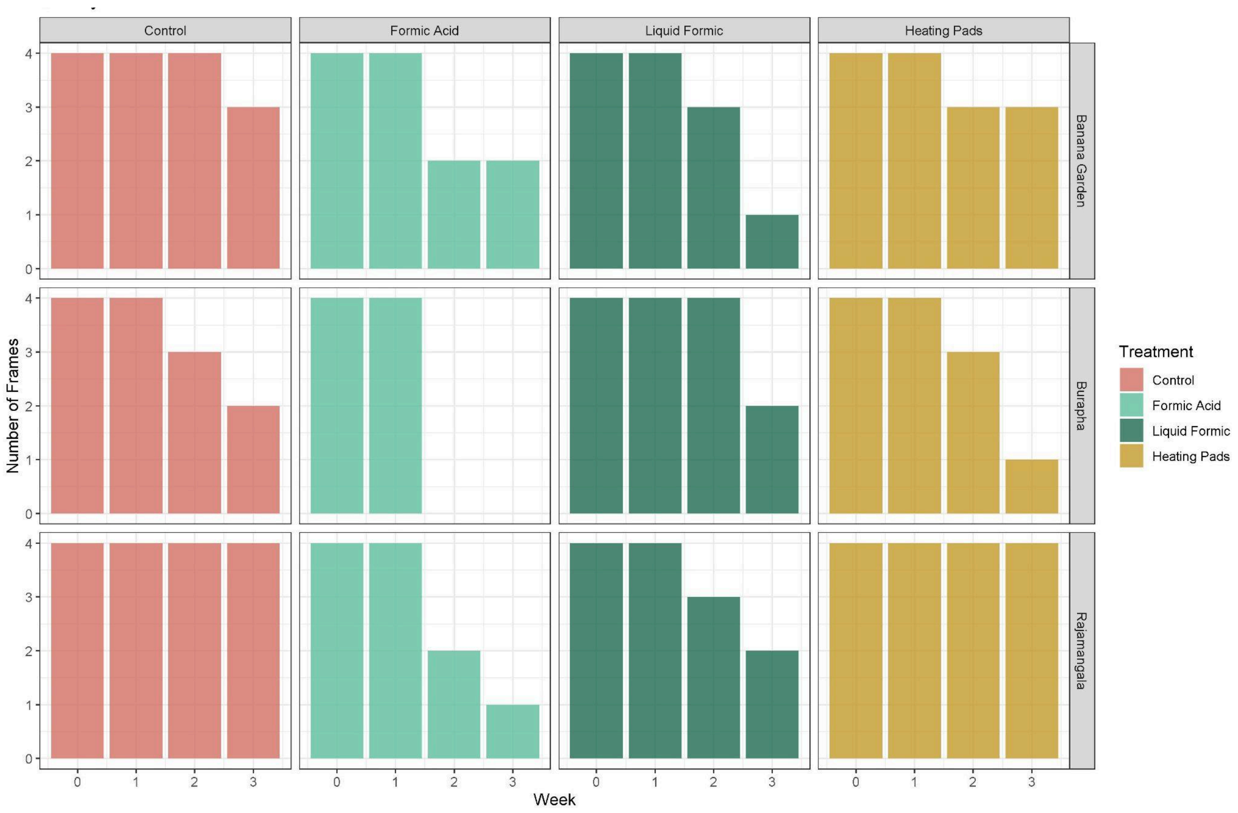


Figure S6. Colony attrition over time. Number of frames that survived throughout the three weeks of the trial.
